# Why species richness of plants and herbivorous insects do or do not correlate

**DOI:** 10.1101/2020.04.29.067504

**Authors:** Naoto Shinohara, Takehito Yoshida

## Abstract

Unraveling the determinants of herbivorous insect diversity has been a major challenge in ecology. Despite the strong association between insect and plant species, previous studies conducted in natural systems have shown great variation in the strength of the correlation between their species richness. Such variation can be attributed to the proportion of generalist insect species (generality), though both higher and lower generality may weaken the correlation because 1) generalist insect species are less dependent on the number of plant species and 2) specialist insect species utilize only a part of the total plant species. To explore these opposing effects, we studied plant and herbivorous insect communities in semi-natural grasslands in Japan. Plant–insect interactions were evaluated in a unique way with a particular focus on the staying and herbivory behaviors of insects, which reflect their habitat use as well as host use. We fousnd that generality of insect communities negatively affected the correlation between species richness of plants and insects. However, such negative effect was significant only when the insect species richness was related with the number of plant species interacted with some insect species, instead of with that of total plant species. The results suggest that considering either of the opposing effects of insect generality is insufficient and they should be inclusively interpreted to understand the relationship between plant and insect species richness.

## Introduction

Understanding the factors determining and regulating species diversity has been a major challenge in ecology (Hutchinson 1959) and several studies have focused on herbivorous insect species, which account for a major fraction of global biodiversity (Novotny et al. 2006; Lewinsohn and Roslin 2008). Due to the strong dependence of herbivorous insects on plants as resources and habitats, herbivorous insect species diversity has been commonly related to plant species diversity (Southwood et al. 1979; Erwin 1982). As expected, studies in experimental grasslands, where plant species richness was manipulated, consistently demonstrated a significant correlation between plant and insect species richness (Haddad et al. 2009; Hertzog et al. 2016). However, contrary to the findings of the experiments, observation studies in natural systems have shown inconsistent patterns about the correlation. Some studies concluded that plant species richness is a good predictor of insect species richness even after accounting for the effects of environmental conditions (Pellissier et al. 2013; Kemp and Ellis 2017; Miller et al. 2017), while others concluded that that was not the case (Hawkins and Porter 2003; Axmacher et al. 2009). Therefore, the relevance of plant species richness to that of insect species is still debatable (Vessby et al. 2002; Castagneyrol and Jactel. 2012).

The variation in the correlation between plant and insect species richness can be attributed to the proportion of generalist insect species (generality), though its effect has been interpreted in opposing directions. As discussed in some studies (Petermann et al. 2010; Kemp and Ellis 2017), lower generality of insect species may weaken the correlation because their limited diet breadth would result in larger part of plant species having no relationship with any insect species. In contrast, other studies (Koricheva et al. 2000; Axmacher et al. 2009; de Araujo et al. 2013) pointed out that a higher proportion of generalist insect species can lead to a weaker correlation because generalist insect species are less dependent on the number of plant species. Although these opposing effects are straightforward, they have been rarely inclusively discussed in previous studies. Therefore, it remains unclear whether higher insect generality dampens or strengthens the correlation of plant and insect species richness in natural systems and few empirical tests were conducted.

To explore the effects of herbivorous insect generality on the correlation between plant and insect species richness, we tested the three hypotheses: (1) if a higher proportion of plant species are not utilized by any insect species, the correlation would be weaker (unutilized plant species hypothesis). This predicts that the herbivorous insect species richness is better explained by utilized plant species richness than by the total plant species richness. (2) Higher proportion of generalist insect species would result in the weaker correlation (higher generality hypothesis), (3) but such pattern would be observed only when the utilized plant species was considered because of the opposing effects of the generality. These hypotheses were tested in a study on plant-herbivorous insect communities in Japanese semi-natural grasslands.

## Materials and Methods

### Study site

The study was conducted in an agricultural area (about 4 km^2^, Fig. S1) located in suburban Tokyo, central Japan (35°35’39.6”N, 139°25’50.9”E). This area has a temperate climate with a mean annual temperature of 15.0 °C, and a mean annual precipitation of 1529.7 mm (data averaged across 1981–2010). The landscape represents the traditional Japanese agricultural landscapes, which comprises a mosaic ensemble of agricultural landscapes, secondary forests, and residential areas in a limited area (Kadoya and Washitani 2011). In this area, semi-natural grasslands are maintained in the form of strips on the edges of rice fields (Fig. S2) by periodic mowing. Although such semi-natural grasslands harbor a great number of plant and insect species (Kitazawa and Ohsawa 2002), changes in local agricultural management (e.g., mowing frequency) affect the diversity of plant and herbivorous insect species (Uchida and Ushimaru 2014). Therefore, a great variation in plant and herbivorous insect species diversity can be observed even within a limited area.

Among this area, 19 transects (2 m × 20 m) were established on the grasslands (Fig. S1, referred to as “sites” hereafter) for the plant and herbivorous insect surveys.

### Data collection

Each site was surveyed in three seasons when plants and herbivorous insects were abundant: early summer (15 June–11 August); late summer (22 August– 5 September); and autumn (19 September –5 October) in 2017. We could not survey every site for every season, because mowing by local farmers often reduced the aboveground vegetation. Therefore, the surveys were restricted to the sites where the vegetational structure seemingly recovered enough after mowing (at least two weeks after mowing) in each season, resulting in 31 surveys in total (9 sites in early summer, 14 sites in late summer, and 8 sites in autumn).

We conducted two types of survey. First, to obtain plant–insect interactions data, we walked along the transect slowly (ca. 20 m/h), while carefully and exhaustively counting the number of herbivorous insect individuals exhibiting either of the two types of the behavior: “herbivory” or “staying”. The behavior was deemed “herbivory” when we observed an insect individual actually feeding on a part of the plant body (i.e., leaf, stem, or petal). The behavior was deemed “staying” when the insect individual just stayed on a part of the plant body. Therefore, the observations of these behaviors were recorded as the interaction data (i.e. what insect species fed or stayed on what plant species). The surveys were conducted twice for each site within each season, resulting in the 2 h sampling effort for each site and each season. Second, to quantify the plant species composition, we established five quadrats (0.5 m × 0.5 m) on each transect and recorded the present plant species with their coverage percentage as an index of abundance.

### Evaluation of plant–insect interactions

Plant–insect interactions were determined by two approaches based on the data collected from the behavior observation. First, we regarded a plant species to be utilized as host by an insect species if “herbivory” behavior was observed between them in any of the transect surveys. Second, we consider non-trophic plant–insect interactions, because architectural structure of plant individuals has been known as an important determinant of insect communities (Woodcock et al. 2007): Some insect species may prefer open microhabitats provided by tall grass species to facilitate thermoregulation (e.g. Joern 2005) and some may prefer particular plant species that attract less predators (“enemy free space hypothesis”; e.g. Price et al. 1980). To evaluate such non-trophic plant–insect interactions, we examined “preferred stays” with an assumption that insect species are dependent on plant species on which they preferably stay. Under this assumption, “preferred stays” of each insect species were determined by comparing “null stays” and observed “staying” behavior. First, the frequency of “null stays” of each insect species on each plant species were evaluated for each season separately. The total (across all sites in each season) number of observed individuals of an insect species were randomly distributed to all plant species proportional to their coverage summed across all sites surveyed in the season. These “null stays” represented the expected frequency of staying of the insect species on each plant species without any preference for plant species. The random distribution procedure was repeated 1,000 times for each insect species, to provide its frequency of “null stays” on each plant species with a 95% confidence interval. Then, the frequency of observed stays on each plant species (unfilled points in Fig. 1) was compared to that of “null stays” and whether it significantly deviated from the expectation (i.e., greater than the upper bound of 95% confidence interval shown as a solid line in Fig. 1) was examined. The plant species, on which focal insect species stayed more frequently than expected (filled points in Fig. 1), were considered as interacted with the insect species. The procedure was conducted for each insect species, and for each season. Although it is interesting to investigate the congruence of trophic and non-trophic interactions, such analysis is beyond the scope of this study and we determined plant–insect interactions by summing up the interaction data obtained from both of the observational and null-model approaches.

**Figure 1.**
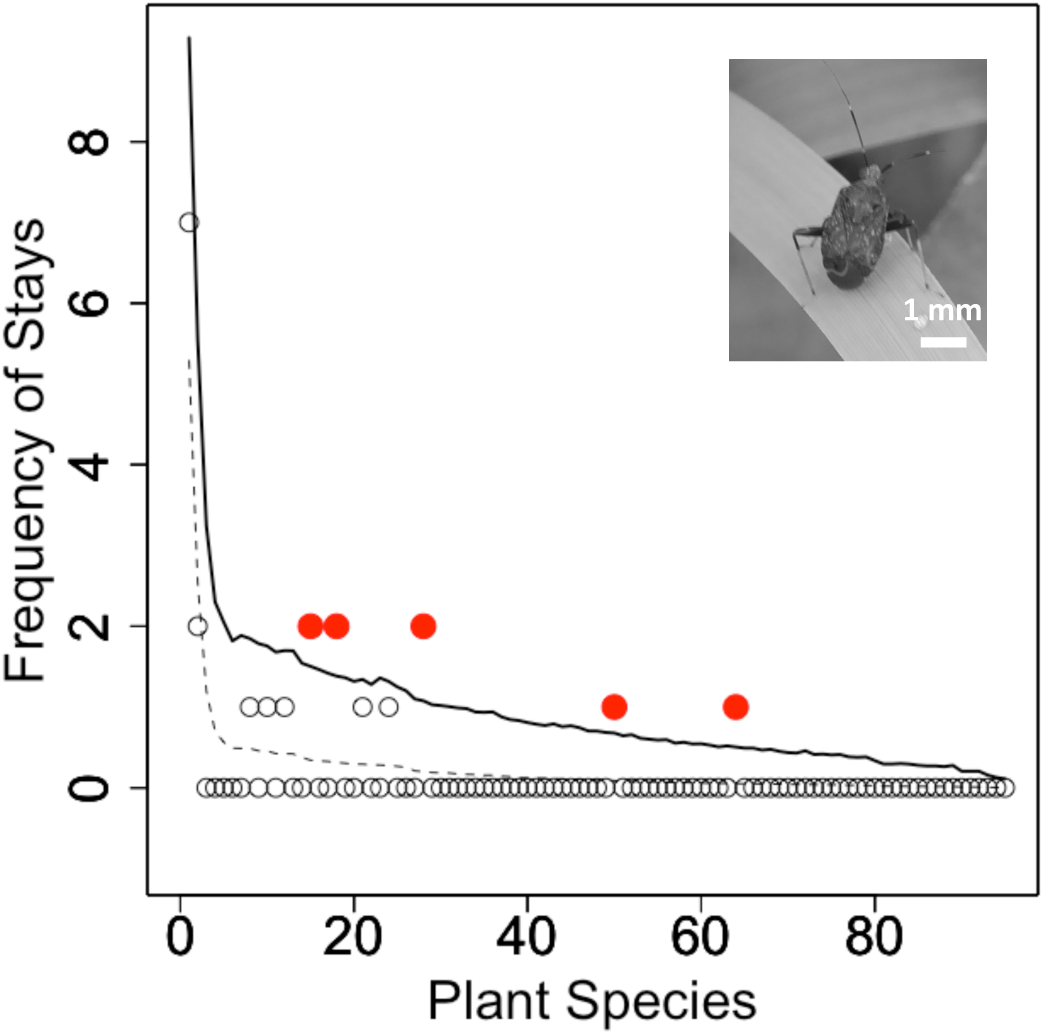
An illustration of the null-model approach for evaluating the preferred stays of a herbivorous insect species (*Proboscidocoris varicornis*), as an example, on all plant species observed. The horizontal axis represents plant species in order of the coverage summed across all sites in each season, and the vertical axis represents the frequency of stays of the herbivorous insect species on each plant species. Dashed and solid lines represent the expected (null) frequency of stays (mean and the upper bound of 95% confidence interval, respectively) calculated based on the total coverage of each plant species summed across all sites in each season. Points represent the frequency of observed stays on each plant species. Filled red points represent that they significantly deviated from the expectation (above the solid line), which we considered the plant and the herbivorous insect species having interacted.

### Statistical analyses

All statistical analyses were performed using R 3.3.1 (R core team 2018). First, diet breadth of each insect species was defined as the number of plant species it interacted with. To represent the proportion of generalist insect species at site (community) level, we calculated the “generality” index that is the averaged diet breadth across insect species within the site.

To test the hypotheses, generalized linear mixed models (GLMMs) were performed using “lme4” package (Bates et al. 2015). First, models were developed to test the hypothesis that insect species richness is better explained by the number of interacted plant species, rather than by that of total plant species. We used GLMMs (Poisson error distribution and log link function) with insect species richness as a response variable, either interacted or total plant species richness as an explanatory variable, and seasons as a random term. Here, the interacted plant species refer to those that were evaluated as interacted with at least one herbivorous insect species within each site.

Second, models were developed to test the second and third hypotheses. We used GLMMs (Poisson error distribution and log link function) with insect species richness as a response variable, either interacted or total plant species richness and its interactive term with generality index as explanatory variables, and season as a random term. The second and third hypotheses predict a negative coefficient for the interactive term, but only when interacted plant species richness was used as an explanatory variable. For the models, the significance of the partial regression coefficients of the explanatory variables were evaluated using Wald test.

## Results

In total, we recorded 152 plant species and 116 herbivorous insect species (1803 individuals). We recorded 118 plant species and 70 insect species (375 individuals) in early summer (9 sites); 93 plant species and 58 insect species (788 individuals) in late summer (14 sites); 75 plant species and 52 insect species (640 individuals) in autumn (8 sites).

We obtained 144 and 1659 observations of herbivory and staying behavior respectively from the field survey. These observations resulted in the estimate of 388 interactions between plant and insect species, where 72 interactions were estimated from observation of herbivory and 328 were from the evaluation of preferred stays, and 12 were from both (Fig. 2a). Distributions of the number of interactions per insect species (Fig. 2b, c) were similar as the one previously known for niche breadth of herbivorous insects, such that specialist species predominate and generalist species are less common (Forister et al .2015).

**Figure 2.**
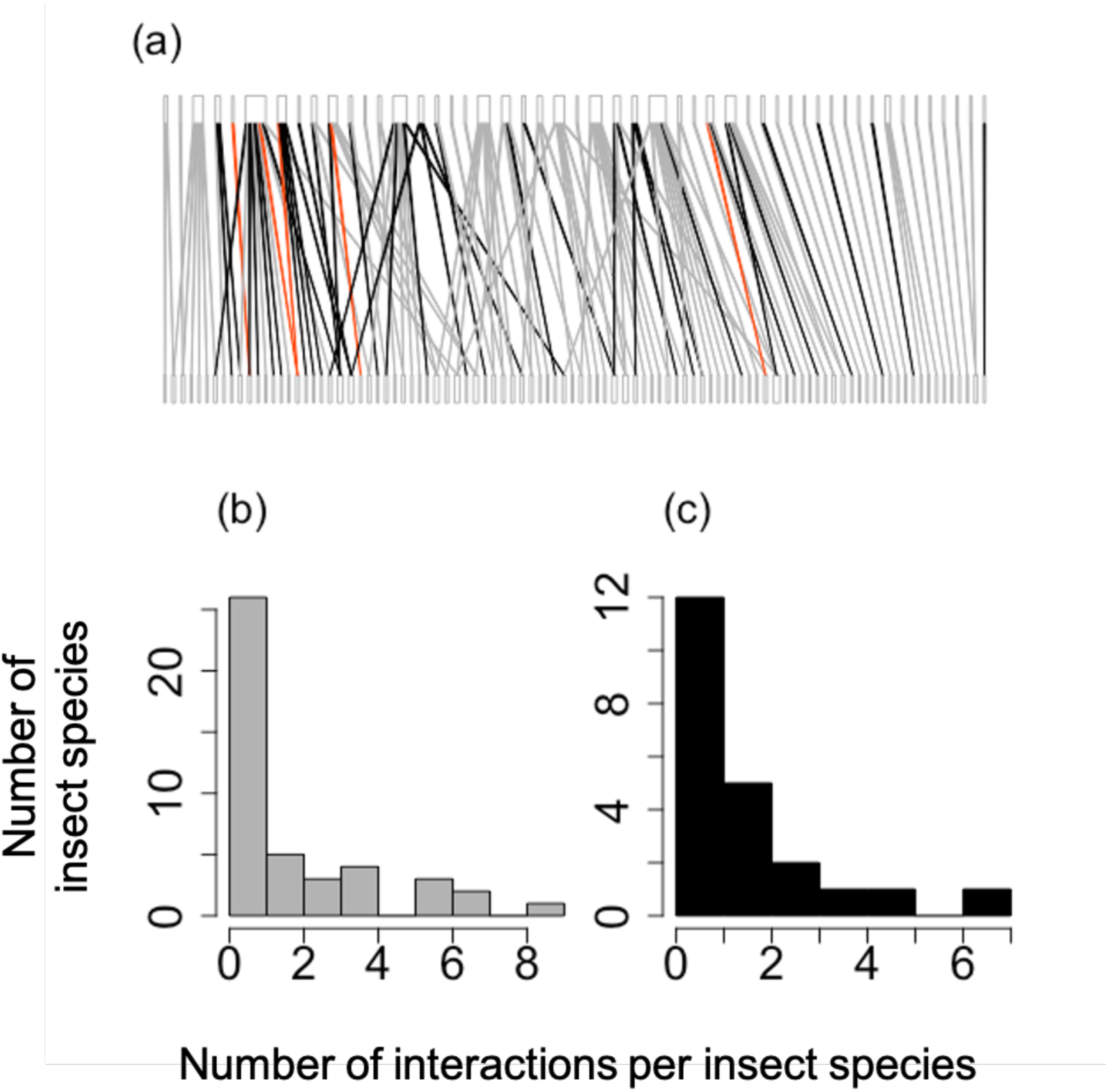
A plant–herbivorous insect interaction network determined by the observational and null-model approaches. Upper and lower bars represent insect and plant species respectively, with their relative frequency of interactions shown by the bar width. Links represent the plant–insect interactions determined by observation of herbivory (black), evaluation of preferred stays (gray), or both (red). The network shown is based on early-summer data as an example (a). Distribution of number of interacted plant species for each insect species, in which interactions were determined by observation of herbivory (b) or evaluation of preferred stays (c).

We found that insect species richness was significantly explained by interacted plant species richness (GLMM: coefficient = 0.044, z = 3.20, P = 0.001) (Fig. 3a), but not by total plant species richness (coefficient = 0.007, z = 0.978, P = 0.328) (Fig. 3b). The GLMM including the interactive term between interacted plant species richness and generality index as an explanatory variable showed a marginally significant negative effect of the interactive term (coefficient = -0.012, z = -1.76, P = 0.078), as well as a significant positive effect of interacted plant species richness (coefficient = 0.089, z = 3.12, P = 0.002) (Fig. 4b). In contrast, when total plant species richness was incorporated into the model, no significant effect of the interactive term (coefficient = -0.001, z = -1.37, P = 0.170) and marginally significant positive effect of total plant species richness (coefficient = 0.024, z = 1.69, P = 0.091) (Fig. 4a) on insect species richness were found.

**Figure 3.**
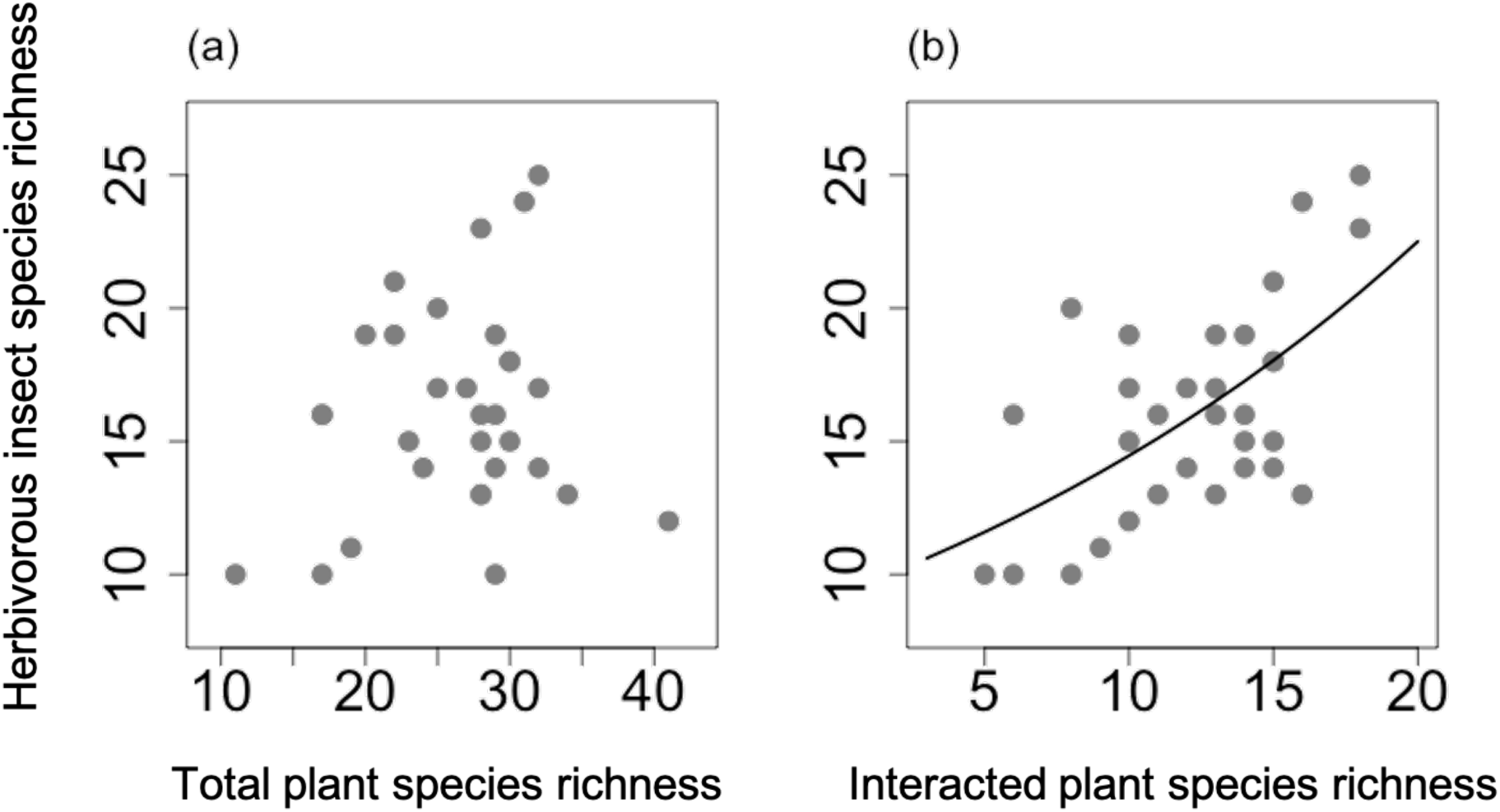
Relationship between herbivorous insect species richness and total plant species richness (a), and relationship between herbivorous insect species richness and interacted plant species richness (b). Regression line is estimated based on the results of the GLMM.

**Figure 4.**
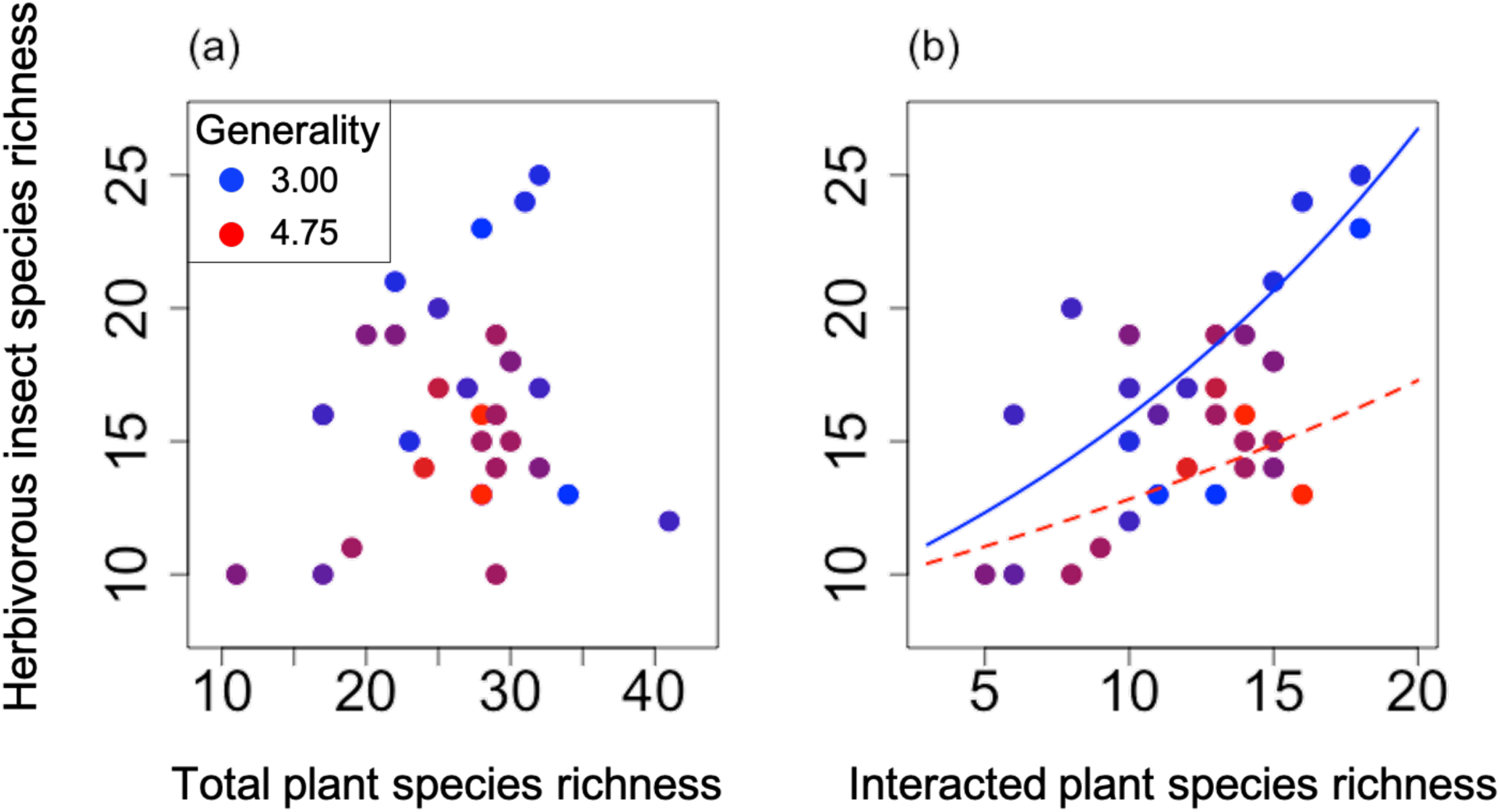
Effects of herbivorous insect generality on the relationships between herbivorous insect species richness and total plant species richness (a), or interacted plant species richness (b). The color of the points represents the generality: reddish ones represent higher generality, and blueish ones represent lower generality. Regression lines are estimated based on the results of the GLMM. Red, dashed line represents the model with the highest generality (4.75), and blue, solid line represents one with the lowest generality (3.00).

## Discussion

Although experimental studies consistently reported that herbivorous species richness is well explained by plant species richness (Haddad et al. 2009; Hertzog et al. 2016), such correlation in natural systems is variable (e.g. Hawkins and Porter 2003; Kemp and Ellis 2017) and consistent explanation for the variation has been lacked. In the present study, we showed that the correlation was dampened by (1) the presence of plant species those were not utilized by any insect species (unutilized plant species hypothesis) and by (2) the high ratio of generalist insect species (higher generality hypothesis), though (3) such effect of higher generality was significant only when the interacted plant species were considered.

In light of the two hypotheses, we may explain the variation in the correlation between plant–insect species richness among previous studies. First, most previous studies have attempted to explain insect species richness by total plant species richness rather than by utilized plant species richness, and some of them have found no significant relationships between them (Vessby et al. 2002; Axmacher et al. 2009; Walcher et al. 2017). On the contrary, a study on plant and their associated insect species (collected from individuals of the focal plants) showed a significant relationship between their species richness even accounting for geographic and environmental variables (Kemp and Ellis 2017). Considering the limited diet breadth of herbivorous insect species (Fig. 2, Forister et al. 2015), distinguishing host plant species would be crucial for explaining insect species richness (but see Hawkins and Porter 2003). The proportion of unutilized plant species depends on the factors causing absence of some insect species independently of host plant species presence (e.g. dispersal limitation or top predator pressure [Koricheva et al. 2000; Schmitz and Suttle 2000]), and such factors may have caused the variation of the correlation between plant–insect species richness among different study systems.

Second, although the negative effect of insect generality on plant–insect species richness correlation may be frequently presumed, empirical evidence that explicitly showed the predicted pattern has been rare. We showed that the correlation between plant and herbivorous insect species richness was negatively affected by insect generality, which indicates that the possibility of colonization of specialist insect species was enhanced with increasing plant species richness, whereas generalist insect species occurred irrespective of plant species richness. It has been suggested that insect generality may vary among study systems that differ in time after disturbance (Piechnik et al. 2008), degree of land-use intensification (de Araujo et al. 2015, Shinohara et al. 2019), spatial scale (Hughes 2000), and taxonomical groups (Forister et al. 2015). Therefore, incorporating those factors in future studies may help explain the relationship between plant–insect species richness. For example, a meta-analysis approach to relate those factors with the variation in the correlation between plant and insect species richness among previous studies may be promising (e.g. Castagneyrol and Jactel. 2012).

More importantly, we found that such negative effect of insect generality on the correlation was not significant when unutilized plant species were also counted in addition to utilized plant species. So far, previous studies have appreciated only either of the opposing directions of the effect of insect generality on plant–insect relationship (unutilized plant species hypothesis and higher generality hypothesis), which has caused the contradictory situation where the weak correlation between plant and herbivorous insect can be associated with both lower (e.g. Kemp and Ellis 2017) and higher (e.g. Axmacher et al. 2009) generality of insects. However, our result suggests that consideration for only negative effect of insect generality may be insufficient to explain the correlation between plant and insect species richness. We conclude that the both mechanisms should be inclusively examined when attributing the strength of the correlation between plant–insect species richness to insect generality.

## Data accessibility

The data used in this study are archived in figshare repository: https://doi.org/10.6084/m9.figshare.11876055.v1

## Acknowledgements

The study was supported by the Research Institute for Humanity and Nature (RIHN: a constituent member of NIHU) Project No. 14200103. We thank Koichi Tagoku, Susumu Yamada, and members of Yui-no-Sato for supporting and assisting our study in the field, and Gaku Takimoto, Marc Johnson, and Yuya Fukano for their helpful comments. We also thank Naoyuki Nakahama, Kazuhide Nakajima, and Takuya Yoshida for assisting field surveys and taxonomic identification.

