## Supplement for "Why species richness of plants and herbivorous insects do or do not correlate"

Figure S1: Location of the study sites in Machida City, Tokyo, Japan. Background aerial photo was taken by WV02 of DigitalGlobe on August 31, 2012.

Figure S2: Photos of the representative study sites. Semi-natural grasslands are maintained in the form of a strip on the edges of rice fields, and the line transects were established on them.

書式を変更: フォント : (英) Times New Roman

移動 (挿入) [1]

Figure S1

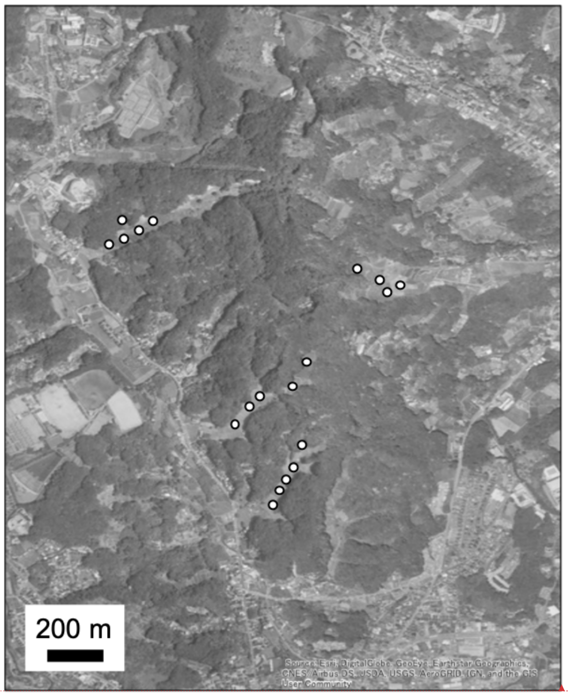


書式を変更: フォント : (英) Times New Roman

書式変更: インデント : 左 : 0 mm, ぶら下げインデント : 5.8 字, 最初の行 : -5.8 字, 行間 : 2 行

書式を変更: フォント : (英) Times New Roman

Figure S2

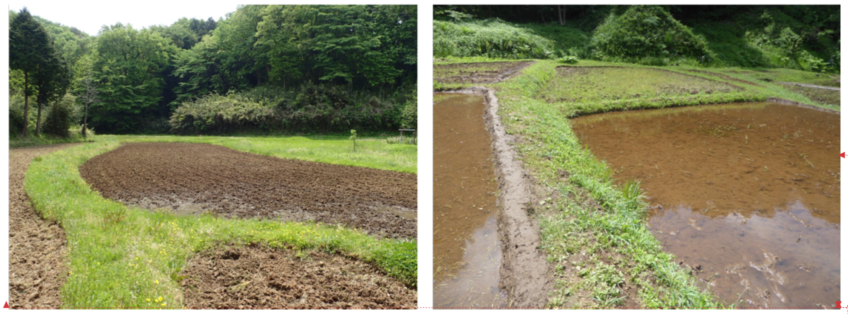

上へ移動 [1]: Figure S2: Photos of the representative study sites. Semi-natural grasslands are maintained in the form of a strip on the edges of rice fields, and the line transects were established on them.

書式を変更: フォント : (英) Times New Roman

書式を変更: フォント : (英) Times New Roman

書式変更: インデント : 左 : 0 mm, ぶら下げインデント : 5.8 字, 最初の行 : -5.8 字, 行間 : 2 行

書式を変更: フォント : (英) Times New Roman

書式変更: 行間 : 2 行

削除:

改ページ

Figure S3: Relationship between the number of interacted plant species, which were evaluated from either of the two approaches (observation of herbivory behavior and evaluation of preferred stays) as interacted with at least one herbivorous insect species in a site, and the number of plant species with eaten marks on their leaves. The dashed line represents the 1:1 relationship.

*<オブジェクト>*

書式を変更: フォント : (英) Times New Roman
